# Csmd2 is a Synaptic Transmembrane Protein that Interacts with PSD-95

**DOI:** 10.1101/362657

**Authors:** Mark A Gutierrez, Brett E Dwyer, Santos J Franco

## Abstract

Mutations and copy number variants of the Cub and Sushi Multiple Domains 2 (*CSMD2*) gene are associated with schizophrenia and autism spectrum disorder. CSMD2 is a single-pass transmembrane protein with a large extracellular domain comprising repeats of Cub and Sushi domains. Although the biological functions of CSMD2 have not been studied, the association between *CSMD2* variants and cognitive function suggest that it may have a role in brain development or function. In this study, we show that mouse *Csmd2* is expressed in excitatory and inhibitory neurons in the brain. Csmd2 protein exhibits a somatodendritic localization in the neocortex and hippocampus, with smaller puncta localizing further out in the neuropil. We show that many of these Csmd2 puncta co-localize with the synaptic protein PSD-95. Using immunohistochemical and biochemical methods, we further demonstrate that Csmd2 localizes to dendritic spines and is enriched in the postsynaptic density. We also find Csmd2 at ribbon synapses of the inner plexiform layer of the retina, suggesting a broader synaptic function of Csmd2 in the central nervous system. Finally, we show that the cytoplasmic tail domain of Csmd2 interacts with synaptic scaffolding proteins of the membrane-associated guanylate kinase (MAGUK) family. The association between Csmd2 and MAGUK member PSD-95 is dependent on a PDZ-binding domain on the Csmd2 tail, which is also required for synaptic targeting of Csmd2. Together, these results point toward a function for Csmd2 in dendrites and synapses, which may account for its association with several psychiatric disorders.

## Introduction

Neurological disorders such as schizophrenia, autism spectrum disorder (ASD), and Alzheimer’s Disease are characterized by deficits in cognitive and social abilities that significantly affect an individual’s quality of life. It is widely hypothesized that these disorders are the result of defects in the capacity of neurons to establish proper connectivity within neural circuits. Most notably, these defects are observed in the contexts of neuronal migration, dendrite development, and synapse formation in the developing cerebral cortex[1, 2]. Such defects would affect the functionality of the neural circuits that give rise to an individual’s higher-order cognitive functions, such as learning and memory. However, the molecular mechanisms that lead to the onset of cognitive disorders remain to be fully understood.

A number of association studies focusing on copy number variants and single nucleotide polymorphisms have identified various novel risk factors for psychiatric disorders. Deletions in members of the Cub and Sushi Multiple Domains (*CSMD*) gene family have been implicated in the occurrence of ASD, schizophrenia, and other neurodevelopmental disorders associated with deficits in cognitive ability and alterations in behavior[3-7]. While recent studies on the *CSMD* family focus on *CSMD1*, little is known about the other members of the family, *CSMD2* and *CSMD3*. *CSMD2* copy number variation is associated with schizophrenia[8], suggesting that *CSMD2* has a significant role in the forebrain. However, the expression patterns, subcellular localizations and functions of CSMD2 in the brain have been previously unknown.

Here, we characterize the biochemical associations and subcellular localizations of Csmd2 in the mouse forebrain. We found that *Csmd2* mRNA and Csmd2 protein are expressed in the mouse neocortex, hippocampus, and retina. Additionally, we provide evidence that Csmd2 is enriched in excitatory and inhibitory neurons in the neocortex and hippocampus. Membrane fractionations of mouse brain homogenates indicate that Csmd2 is enriched in synaptosome-containing fractions, particularly in the postsynaptic density (PSD). We have validated these findings via immunohistochemistry, showing that Csmd2 puncta localize to dendritic spines and colocalize with the postsynaptic scaffold protein PSD95. Utilizing yeast 2-hybrid screening as well as co-immunoprecipitation assays, we find that the intracellular tail domain of Csmd2 interacts with PSD95. This interaction depends on the PDZ-binding motif on Csmd2, and mutation of this PDZ ligand abolishes the interaction with PSD95 and the synaptic enrichment. Taken together, these results indicate that Csmd2 is a transmembrane protein localized at synapse in the central nervous system, suggesting a role in synaptic function that may be perturbed in certain neuropsychiatric disorders.

## Results

### Csmd2 mRNA is expressed in the mouse neocortex and hippocampus

The mouse *Csmd2* gene comprises 71 exons and is predicted to encode a 13,555 base long mRNA (Fig. 1A). While cloning the full-length cDNA from postnatal forebrain, we also identified a splice variant in which exon 7 splices to exon 14 (Fig. 1A). The protein encoded by the full-length mRNA is predicted to be 3,611 amino acids with an approximate molecular weight of 392 kDa. The TMHMM 2.0 Server (http://www.cbs.dtu.dk/services/TMHMM/) [9] predicts a single transmembrane helix in the mouse Csmd2 protein at amino acids 3534-3556 (Fig. 1A). Results from the TatP 1.0 Server (http://www.cbs.dtu.dk/services/TatP/) [10] prediction indicates the presence of a signal peptide in the N-terminal 37 amino acids of Csmd2, with a likely cleavage site between positions 37-38 (Fig. 1A). The large extracellular domain of Csmd2 contains 14 complement C1r/C1s, Uegf, Bmp1 (CUB) domains, each separated by an intervening sushi domain (Fig. 1A). Following the CUB/sushi repeats is a series of 15 consecutive sushi domains, the transmembrane domain and a cytoplasmic tail domain at the C-terminus.

**Figure 1.**
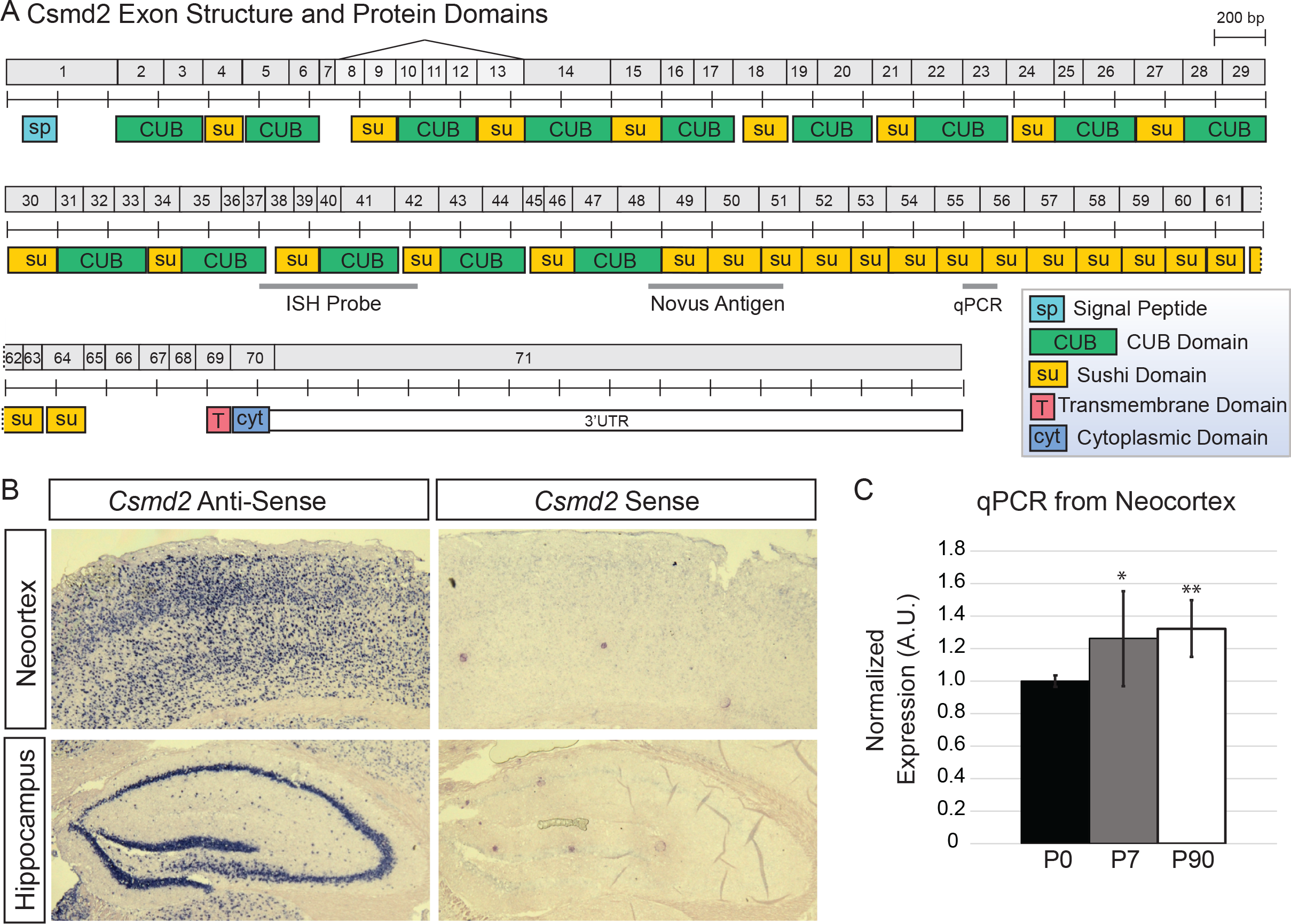
*Csmd2* mRNA is expressed in the mouse forebrain. ***A***, Schematic of numbered exons of mouse *Csmd2* mRNA and domain structure of mouse Csmd2 protein, noting locations of alternative splicing, probes used for in situ hybridization and quantitative PCR analysis, and antigen used to generate the anti-Csmd2 antibody from Novus. ***B***, In situ hybridization visualizes the broad expression of *Csmd2* mRNA throughout all neuronal layers in the neocortex and hippocampus. A sense-strand probe was used as a negative control. ***C***, Quantitative PCR analysis shows a slight increase in *Csmd2* mRNA expression in the neocortex from timepoint P0 to P7 and P90. Values were normalized to *Cyclophilin A* expression and plotted relative to the P0 timepoint. * P < 0.05, ** P < 0.001.

Publically available databases show that human *CSMD2* mRNA (https://www.proteinatlas.org) [11] and mouse *Csmd2* mRNA (http://www.informatics.jax.org/expression.shtml) [12] expression is highest in the central nervous system. To futher analyze *Csmd2* mRNA expression in the adult mouse forebrain, we performed RNA *in situ* hybridization using a probe spanning exons 37-42 (Fig. 1A). We found *Csmd2* mRNA widely-expressed throughout the neuronal layers of the adult mouse neocortex and hippocampus (Fig. 1B). Quantitative real-time PCR analysis showed that *Csmd2* expression slightly increased in the neocortex during the first postnatal week, at which time it reached similar levels as in the adult (Fig. 1C). Together, these data indicate that mouse Csmd2 is a large, single-pass transmembrane protein expressed in the developing and mature forebrain.

### Csmd2 protein is expressed in neurons in the mouse forebrain

We next wanted to determined Csmd2 protein expression and localization in the forebrain. First, we validated several commercially available anti-Csmd2 antibodies. We generated cDNA expression plasmids for either full-length Csmd2, or a truncated form of Csmd2 in which the ectodomain contains only the 15 Sushi repeats proximal to the transmembrane domain (Csmd2 15x; Fig. 2A). Both constructs included a 3x-FLAG tag at the N-terminus. Upon transfection of these constructs into HEK293T cells, Western blot analysis using 3 different anti-Csmd2 antibodies revealed the predicted 380-kDa band corresponding to full-length Csmd2 only in the transfected conditions (Fig. 2A). Immunoprecipitation of these samples using anti-FLAG beads prior to Western blotting confirmed that the bands seen in each condition corresponded to the exogenous FLAG-tagged Csmd2 protein (Fig. 2A). Only the anti-Csmd2 antibody from Novus was able to detect the truncated Csmd2 15x protein (Fig. 2A), indicating that the other 2 antibodies recognize antigens more N-terminal on Csmd2. We further tested the antibodies by fluorescence immunocytoochemistry on HEK293T cells cotransfected with full-length FLAG-Csmd2 together with the transfection marker myristoylated tdTomato (myrtdTomato). We confirmed that all 3 antibodies labeled plasma membranes only in the transfected HEK293T cells, but not in neighboring untransfected cells (Fig. 2B). Furthermore, we confirmed colocalization of the FLAG tag and Csmd2 in transfected cells using an anti-FLAG antibody (Fig. 2C). These data indicate that all 3 antibodies can detect mouse Csmd2 protein in Western blots and immunocytochemistry.

**Figure 2.**
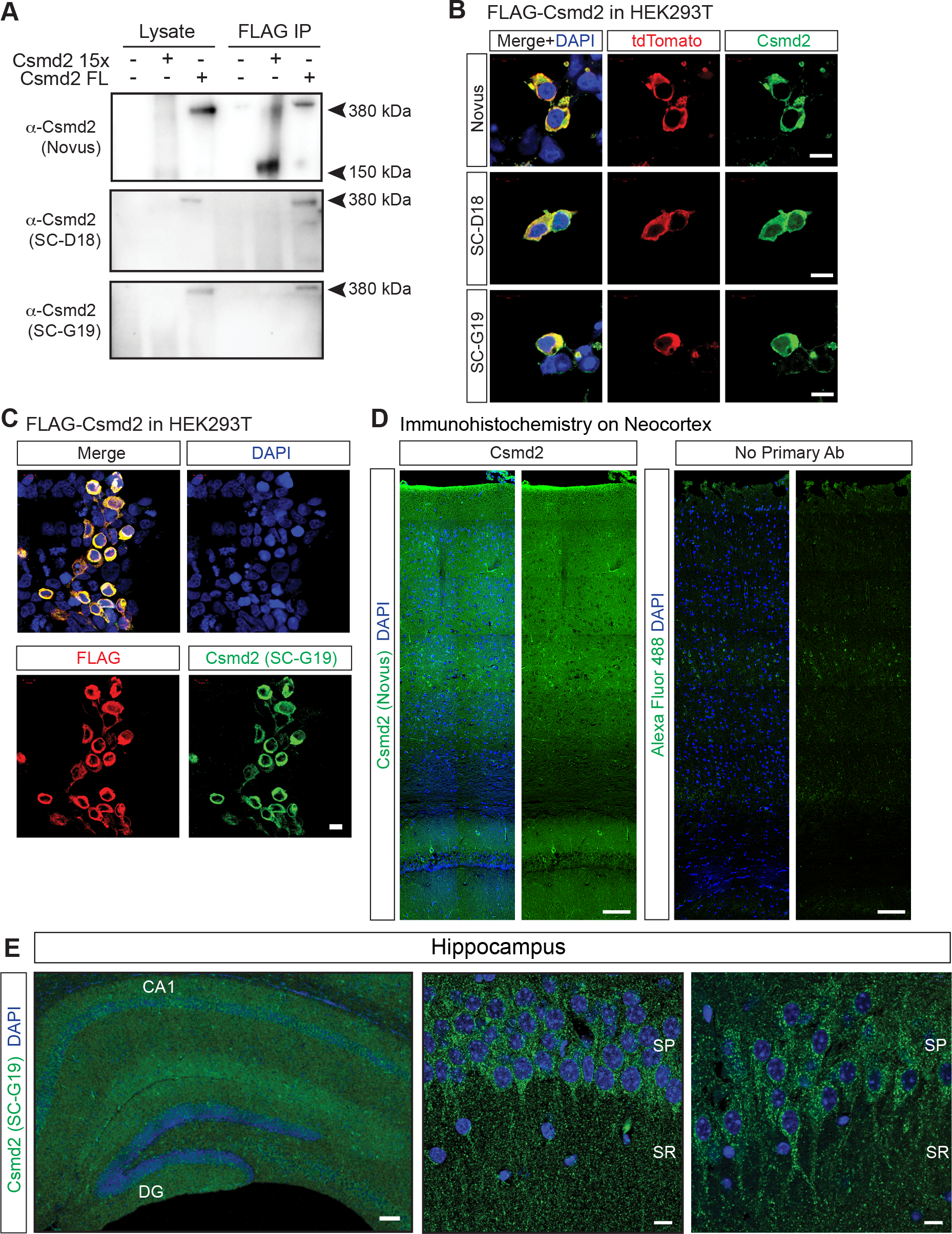
Validation of Csmd2 Antibodies and Detection of Csmd2 protein in the Mouse Forebrain. ***A***, HEK293T cells were transfected with expression constructs for FLAG-tagged full-length (Csmd2 Full-Length) or truncated (Csmd2 15x) Csmd2. The Novus Csmd2 antibody detects both full-length and truncated forms in Western blot analysis. SC-D18 and SC-G19 antibodies only detect the full-length construct. ***B***, Fluorescence immunocytochemistry of HEK293T cells co-transfected with FLAG-Csmd2 full-length and myristoylated tdTomato as a transfection marker. All 3 antibodies recognize overexpressed Csmd2 in the transfected cells. ***C***, As in (B), but stained with anti-FLAG antibody. SC-G19 signal colacalizes with FLAG signal. ***D***, Coronal section of adult mouse neocortex stained for Csmd2 (Novus) shows Csmd2 expression throughout neuronal layers of the neocortex. No signal is seen in the absence of primary antibody (right panels) ***E***, Csmd2 expression in the adult mouse hippocampus appears broad throughout the neuronal layers, as seen in the whole-view image (left). Zoom-in images show somato-dendritic patterns in cell bodies and punctate patterns in the neuropil. Scale bars: B-C, 10 μm; D, 100 μm; E left panel, 100 μm; E zoom right panels, 10 μm. CA1, cornu ammonis 1; DG, dentate gyrus; SP, stratum pyramidale; SR, stratum radiatum.

We conducted fluorescence immunohistochemistry on coronal sections from P90 mouse brains to determine Csmd2 protein localization in the mouse forebrain. We observed Csmd2 signal distributed throughout the neocortex and hippocampus. In the neocortex, Csmd2 was expressed throughout the neuronal layers, similar to our in situ hybridization findings (Fig. 2D). The vast majority of Csmd2 staining in the neocortex was lost in the absence of primary antibody. Similar to the neocortex, Csmd2 was widely-expressed throughout the hippocampal layers (Fig. 2E). Higher magnification images of neurons in the CA1 layer revealed that Csmd2 exhibited a somato-dendritic pattern that extended into the apical dendrites of neurons from cell bodies in the stratum pyramidale. In the stratum radiatum, Csmd2 signal was found in smaller puncta throughout the neuropil. Taken together, these data show that Csmd2 is widely expressed throughout the mouse neocortex and hippocampus, adopting somato-dendritic and punctate expression signals in neurons.

### Csmd2 is expressed in excitatory projection neurons and inhibitory interneurons

To elucidate what cell types express Csmd2, we probed P90 mouse neocortical sections for Csmd2 (SCG19) along with Ctip2 for corticospinal motor neurons, Satb2 for corticocortical projection neurons, PV and SST for interneurons, Olig2 for oligodendrocytes and Aldh1L1 for astrocytes (Fig. 3). We observed Csmd2 expression in Ctip2^+^ and Satb2^+^ cells, demonstrating that Csmd2 is expressed in excitatory projection neurons. Additionally, Csmd2 was expressed more strongly in PV^+^ and SST^+^ cells, indicating higher expression in inhibitory interneurons (Fig. 3). In each of these cases, Csmd2 exhibited a clear somato-dendritic expression pattern, similar to that seen in the hippocampus. Csmd2 was expressed weakly in Aldh1L1^+^ astrocytes and was undetectable in Olig2^+^ oligodendrocytes (Fig. 3). Identical patterns were observed with the other 2 Csmd2 antibodies (data not shown), but the signal-to-noise ratio was not as robust as with the SC-G19 antibody. These data indicate that Csmd2 is expressed by excitatory neurons, inhibitory interneurons, and astrocytes in the mouse forebrain.

**Figure 3.**
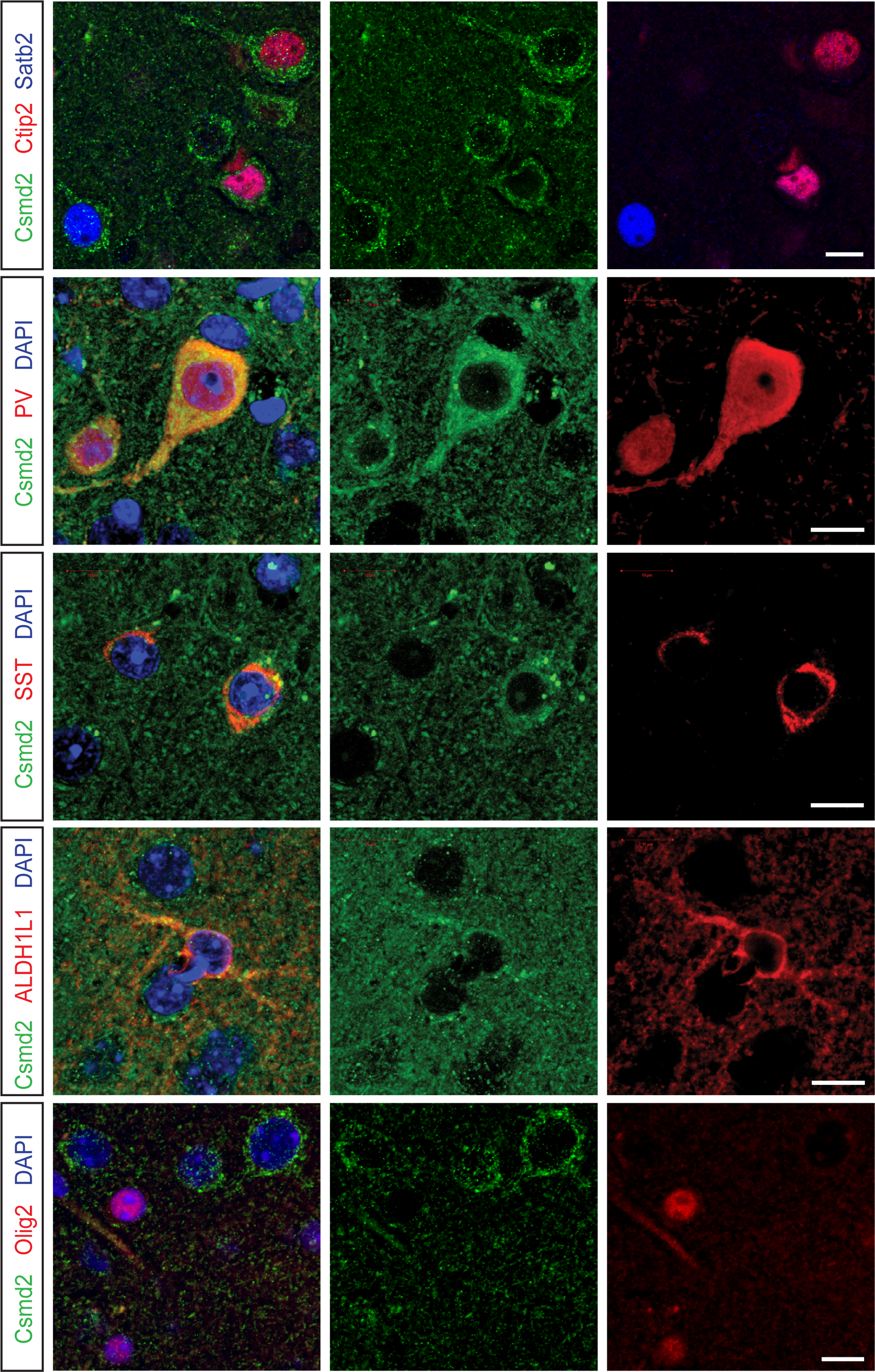
Csmd2 is Expressed in Multiple Cell Types. Coronal sections of P90 mouse neocortex. Fluorescence immunohistochemistry revealed expression of Csmd2 in Ctip2+ and Satb2+ excitatory projection neurons, PV+ and SST+ inhibitory interneurons, and ALDH1L1+ astrocytes. Csmd2 expression was not observed in Olig2+ oligodendrocytes in the white matter or gray matter. Scale bars, 10 μm.

### Csmd2 is enriched in synapses

Association studies have implicated Csmd protein family loss of function in the onset of cognitive decline[3, 5, 7, 8]. This suggests that Csmd2 may function to maintain normal synaptic function in the brain. Supporting this hypothesis is evidence that other Cub- and Sushi-containing proteins such as Lev9, Lev10, Neto1, and Neto2 play a critical role in the regulation of synaptic activity. Specifically, these proteins act as auxiliary subunits that facilitate receptor trafficking, clustering, and clustering of these receptors at the synapse[13-17]. Therefore, we speculated that Csmd2 might play a role at synapses. To begin to test this hypothesis, we used several complementary methods to determine if Csmd2 is localized to synapses in the mouse forebrain. First, we devised a strategy to help visualize individual synapses in vivo. We used *in utero* electroporation to introduce expression plasmids into progenitors of excitatory neurons in the cortex (Fig. 4A). We electroporated a myristoylated tdTomato construct together with an expression plasmid for a GFP-tagged intrabody targeting endogenous PSD-95, allowing us to visualize dendritic spines and postsynaptic densities of excitatory neurons in the mature cortex (Fig. 4A). When combined with Csmd2 IHC, we readily found Csmd2 puncta co-localized with PSD-95 at the ends of dendritic spines (Fig. 4B). Interestingly, we noticed that not all PSD95^+^ spines displayed Csmd2 punctate signals. To follow up this observation, we labeled hippocampal neurons with myrtdTomato in utero at E14.5 and then prepared primary hippocampal neuron cultures at E17.5. Fluorescence immunocytochemistry probing for Csmd2 at 14 days in vitro (14 DIV) revealed that approximately 80% of labeled spines contain some Csmd2 signal (Figure 5A-B). To test for synaptic localization in a third system, we performed immunohistochemistry on P90 mouse retinal sections. We found Csmd2 localized throughout the retinal layers, including in a somatodendritic pattern in the inner nuclear layer and in punctate patterns in both the inner and outer plexiform layers (Fig. 5C). Higher magnification images revealed Csmd2 concentrated at the center of PSD-95+ ribbon synapses in the outer plexiform layer (Fig. 5C).

**Figure 4.**
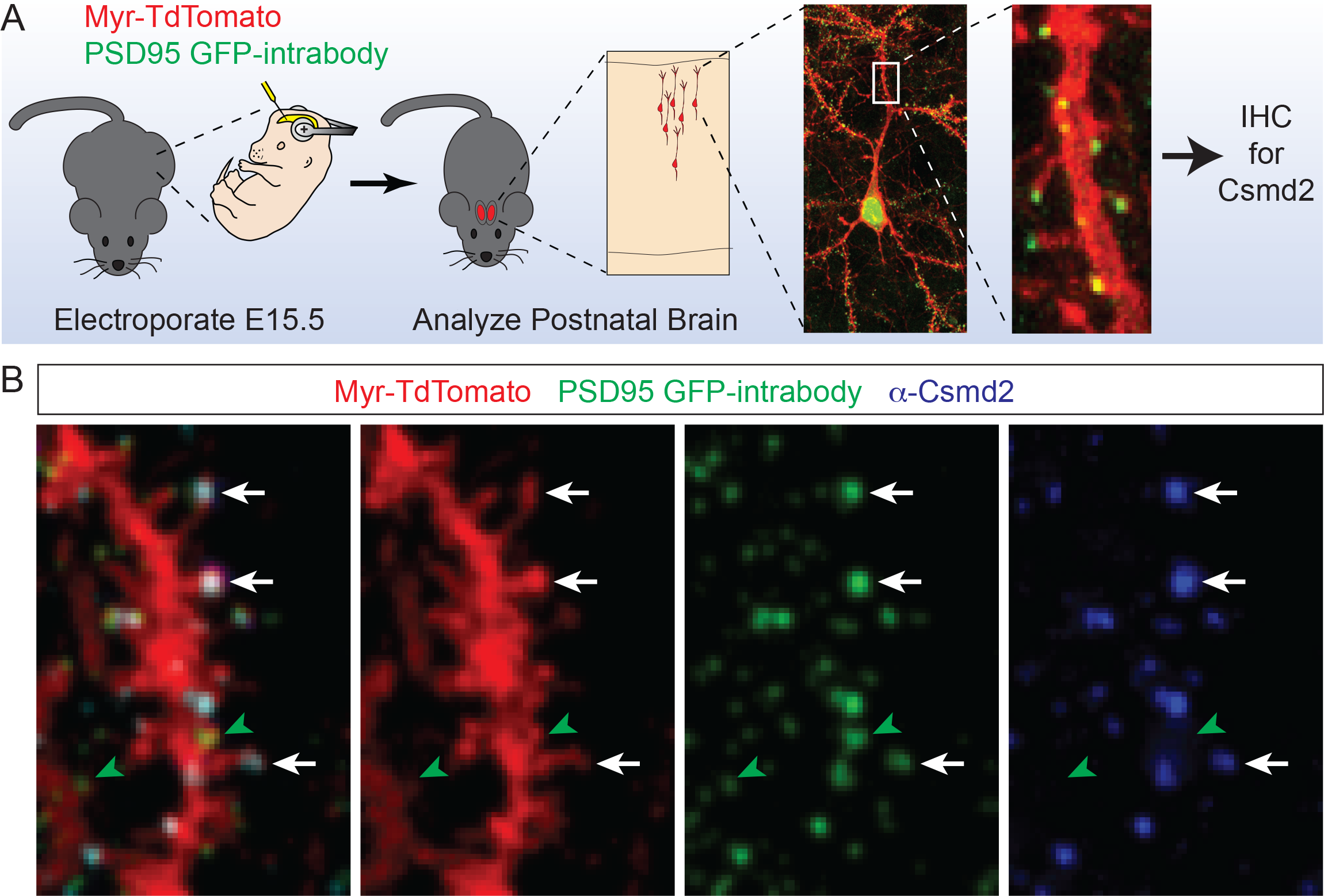
Csmd2 Co-Localizes with PSD95 at Synapses. ***A***, Schematic of experimental approach for in vivo labeling of neuronal dendritic spines with myrisotylated tdTomato and post-synaptic densities with a GFP-fused intrabody targeting PSD-95. Electroporated brains were stained for Csmd2 (SC-G19) at P30. ***B***, Immunohistochemical analysis of P30 neurons after in utero electroporation show localization of punctate Csmd2 at PSD-95^+^ synapses on both the dendritic shaft and at the ends of dendritic spines (white arrows). A subset of PSD-95^+^ puncta are not positive for Csmd2 (green arrowheads).

**Figure 5.**
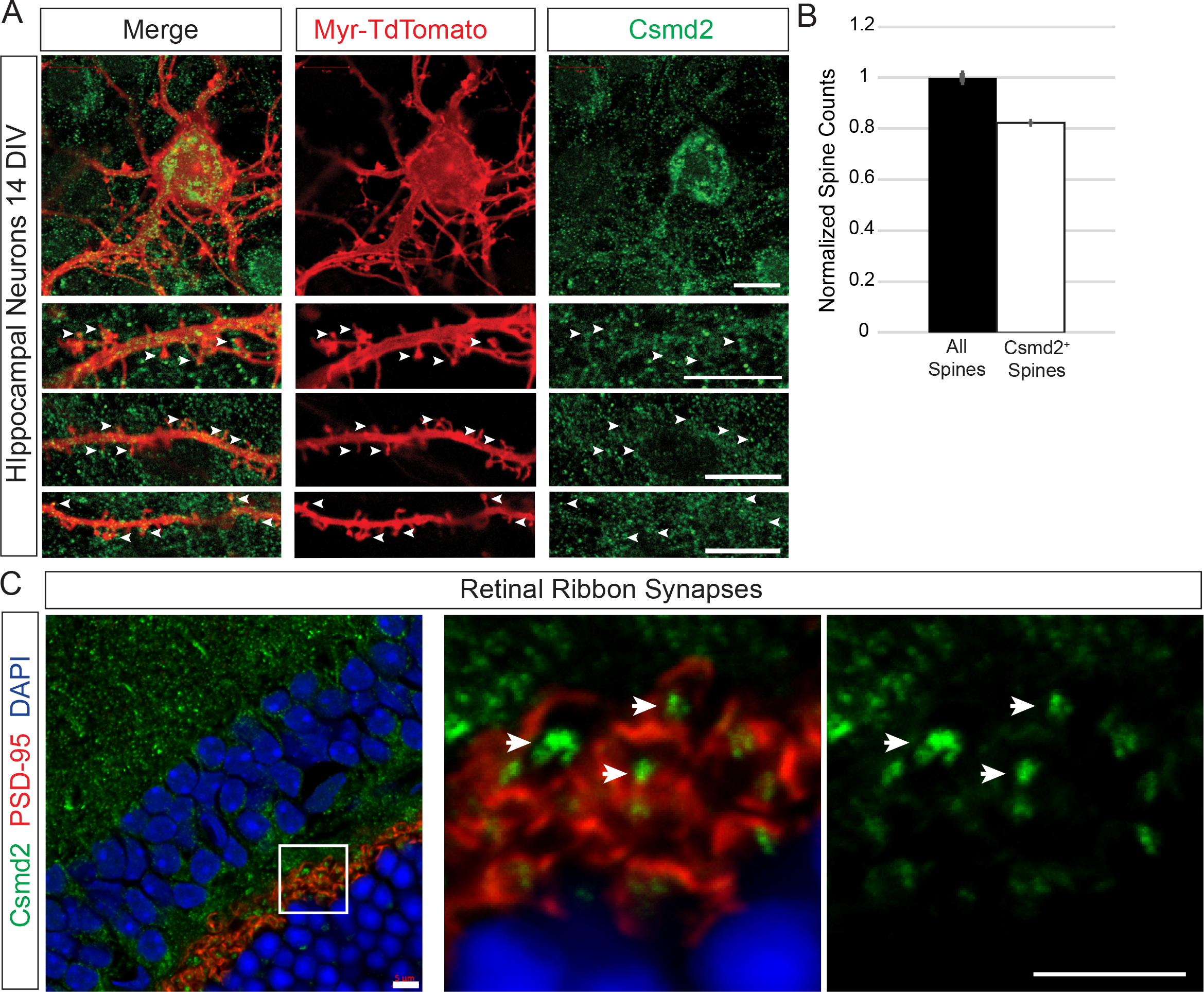
Csmd2 localizes to dendritic spines in vitro and to retinal ribbon synapses. ***A***, 14 DIV hippocampal neurons show somato-dendritic Csmd2 staining (upper panels) and punctate expression throughout their dendrites, including in dendritic spines (lower panels). ***B***, Quantification of Csmd2 punctate expression reveals Csmd2 expression at more than 80% of dendritic spines at 14 DIV. ***C***, Fluorescence immunohistochemistry of P90 mouse retina reveals the expression of punctate Csmd2 at the center of ribbon synapses in the innerplexiform layer. Scale bars: A, 10 μm; C, 5 μm.

To further characterize the subcellular localization of Csmd2 in forebrain neurons, we isolated synaptosomal fractions from P30 mouse whole brain tissue using a Percoll gradient[18], which allows for the separation of small membranes, myelin, membrane vesicles and synaptosomes (Fig. 6A). We ran equal amounts of protein from each fraction on an SDS-PAGE gel. Upon Western blot analysis of the fractions, we observed enrichment of Csmd2 in synaptosome-containing fractions F3 and F4, along with PSD-95 (Fig. 6A). Csmd2 could be detected in syntaptosomal fractions by all Csmd2 3 antibodies. To determine which compartment of the synaptosome Csmd2 was localized, we utilized a second method for the fractionation of the postsynaptic density (PSD) from a crude synaptosomal preparation[19]. We confirmed by this method that Csmd2 was found in the synaptosomal pellet (P2) fraction, specifically in the Triton-X-insoluble PSD pellet fraction (TxP) (Fig. 6B). Taken together, these data show that Csdm2 is localized to synapses in the neocortex, hippocampus and retina.

**Figure 6.**
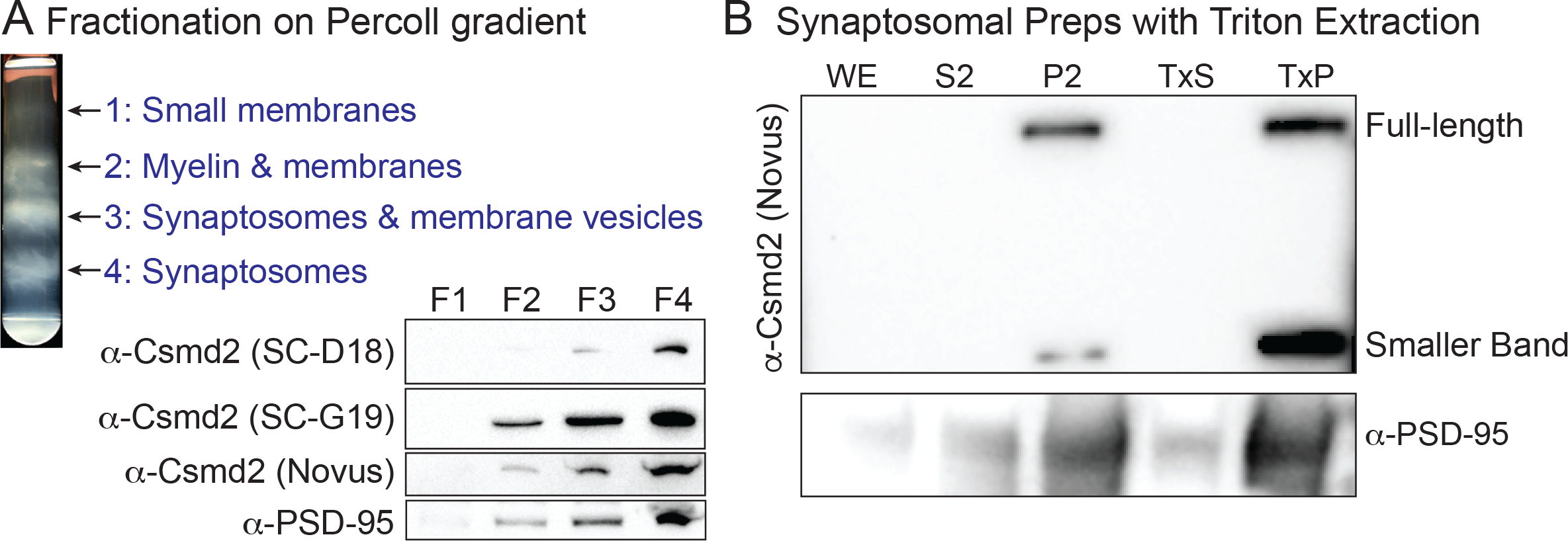
Csmd2 is found in synaptosomal and postsynaptic fractions. ***A***, Membrane fractionation of P30 mouse forebrain lysate using a Percoll gradient. Western blot analysis using all 3 Csmd2 antibodies shows enrichment of Csmd2 and PSD-95 in synapto-some-containing fractions. ***B***, Preparation of crude synaptosomes shows Csmd2 enriched in the synaptosomal fraction (P2) but not the soluble fraction (S2). Csmd2 levels are too low to be detected in the whole extract (WE) at this loading concentration. Preparation of crude post-synaptic densities shows Csmd2 and PSD-95 enriched in the post-synaptic density fraction (TxP) but not in the Triton-soluble fraction (TxS). Triton extraction of synaptosome-containing fractions reveals the enrichment of Csmd2 at the postsynaptic density together with PSD-95. The Csmd2 Novus antibody also recognizes a smaller band at approximately 150 kDa, which may be a cleavage product or alternative splice form. WE, whole extract; S2, supernatant from lysate; P2, crude synaptosomal pellet fraction; TxS, pre-/extra-synaptic soluble fraction; TxP, postsynaptic density pellet fraction.

### Csmd2 interacts with synaptic scaffold proteins

To begin to study the possible functions of Csmd2 in the brain, we wanted to elucidate some of the molecular associations of Csmd2. We employed a yeast 2-hybrid screen using the intracellular portion of Csmd2 as bait protein, and an adult mouse brain library as prey. Our screen identified 7 proteins that interacted with the Csmd2 cytoplasmic tail domain with high or very high confidence (Fig. 7). Interestingly, several of the identified interactors are known synaptic scaffolding proteins of the membrane-associated guanylate kinase (MAGUK) family, including SAP-97, PSD-93, and PSD-95. Each interaction mapped to a specific PDZ domain within the prey. We found that Csmd2 contains a putative class I PDZ-binding motif (TRVCOOH) at the C-terminus of its cytoplasmic tail.

**Figure 7.**
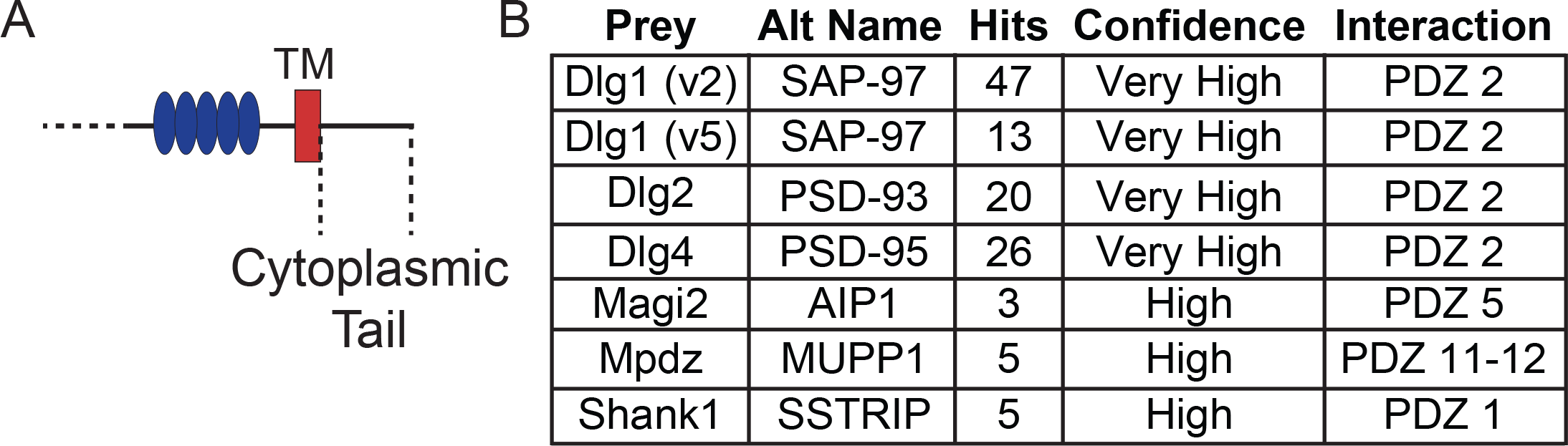
Candidate Csmd2 intracellular interaction partners. ***A***, Yeast 2-hybrid screen using the cytoplasmic tail domain of mouse Csmd2 (schematized on the left) as bait protein and an adult mouse brain library as prey. ***B***, Results of the 2-hybrid screen reveal high-confidence hits with synaptic scaffolding proteins. All interactions were mapped to specific PDZ domains within these multi-PDZ proteins.

As a starting point to validate our 2-hybrid results, we performed immunoprecipitation of PSD-95 from mouse adult brain lysates and found that Csmd2 co-immunoprecipitated with PSD-95 (Fig. 8A). Furthermore, when we co-expressed PSD-95 with the FLAG-tagged Csmd2 construct (Fig. 8B) in HEK293T cells, FLAG-Csmd2 co-immunoprecipitated upon PSD-95 pull-down (Fig. 8C). To determine if the interaction between PSD-95 and Csmd2 is dependent on PDZ/PDZ-ligand interactions [20], we generated a construct in which the Csmd2 PDZ-binding domain was mutated from TRV to AAA (Fig. 8B). The interaction between Csmd2 and PSD-95 was completely abolished when the PDZ-binding motif in Csmd2 was mutated (Fig. 8C). These data confirm that Csmd2 interacts with PSD-95 via a PDZ-binding domain at the C-terminus of the cytoplasmic tail.

**Figure 8.**
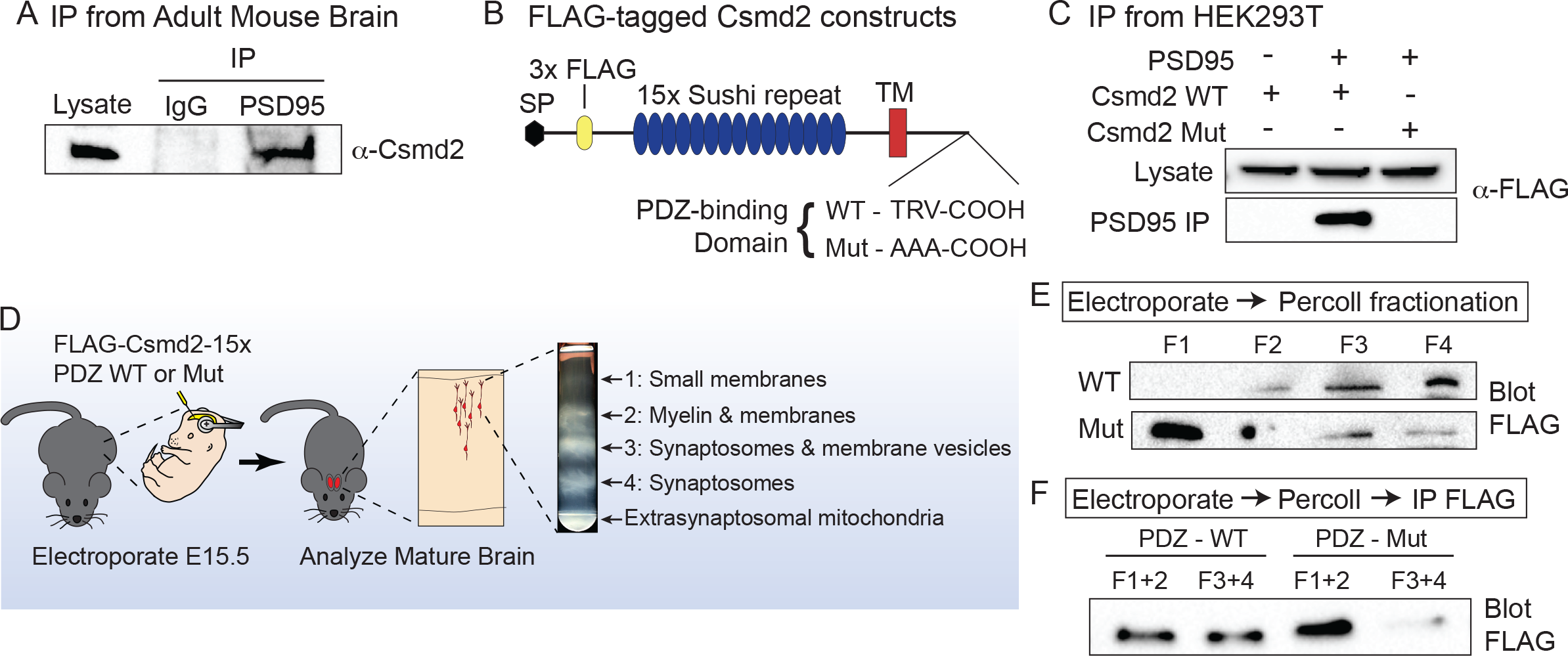
Csmd2 interacts with PSD-95 via a PDZ-binding domain. ***A***, Endogenous Csmd2 from adult mouse brain lysates co-immunoprecipitates with PSD-95, but not IgG control. ***B***, Truncated Csmd2 expression constructs. In Mut, the PDZ-binding motif (TRV) is mutated to AAA. ***C***, Constructs from (C) transfected into HEK293T cells ± PSD-95 cDNA. WT, but not Mut, co-IPs with PSD-95 after anti-PSD-95 pull-down. ***D***, Constructs from (C) were electroporated into the cortex at E15.5 and the electroporated region was microdissected from adult brains and fractionated on a Percoll gradient. ***E***, Equal amounts of protein from each fraction were fun on an SDS-PAGE gel and Western blotted for FLAG. Csmd2 WT is enriched in synaptosomal fractions F3-F4, but Mut is enriched in non-synaptosomal fractions F1-F2. ***F***, Similar experiment as in (E), but fractions were pooled in pairs and immunoprecipitated with α-FLAG before Western blot. Csmd2 Mut is lost from synaptosomal fractions F3-F4.

Based on our data showing colocalization of Csmd2 with PSD-95 at synapses, we hypothesized that the Csmd2 PDZ domain is important for synaptic localization of Csmd2. To test this, we conducted *in utero* electroporation experiments to express wild-type FLAG-Csmd2 or the version in which the PDZ-binding domain is mutated (Fig. 8D). We electroporated wild type or Csmd2 PDZ mutant constructs into embryonic mouse cortices at E15.5 and conducted Percoll fractionations at P30, followed by Western blot analysis (Fig. 8D). Similar to endogenous Csmd2, FLAG-Csmd2 was found enriched in synaptosome-containing fractions (Fig. 8E). Conversely, the PDZ-binding mutant version was primarily found in fractions 1 and 2 (Fig. 8E). Even when we increased sensitivity of the assay by employing a FLAG IP to enrich for the tagged protein, we could barely detect any mutant version in the synaptosomal fractions F3-F4 (Fig. 8F). We concluded that the synaptic localization of Csmd2 depends upon its intracellular PDZ-binding domain, possibly through its interactions with PDZ-containing synaptic scaffold proteins.

## Discussion

Genetic variations in the CSMD genes have been associated with the onset of schizophrenia and ASD in a number of previously-published GWAS studies in a manner suggesting that the reduced expression of the CSMD proteins results in the decline of cognitive function[4-8]. However, a characterization of the Csmd proteins in neuronal cell biology, particularly in the context of connectivity in a neural circuit has yet to be reported. Here, we show that Csmd2 is expressed in the mouse forebrain and localizes to the dendritic spines of neurons through a PDZ/PDZ-ligand interaction with the synaptic scaffold proteins. Immunostaining reveals that Csmd2 expression exhibits defined somato-dendritic and punctate patterns. Additionally, we show that Csmd2 is enriched in Ctip2^+^ and Satb2^+^ excitatory neurons, PV^+^ and SST^+^ interneurons, and ALDH1L1^+^ astrocytes. Mulitple immunohistochemical and biochemical approaches demonstrated the synaptic localization of Csmd2. We further showed that the interaction of Csmd2 with PSD-95 and the synaptic localization of Csmd2 is largely dependent on its PDZ-binding domain in the cytoplasmic tail. Taken together, these data indicate that Csmd2 is a novel synaptic transmembrane protein, point toward a synaptic function for this previously uncharacterized protein.

Although little is known about the functions of Csmd2 in the brain, we may gain some insight from the roles of other Cub and Sushi domain containing proteins. For example, Cub/Sushi-containing proteins such as Lev9/10 and Neto1/2 play significant roles as accessory subunits of synaptic receptors. Specifically, Neto1 and Neto2 are responsible for phosphorylation-dependent regulation of kainate receptor subunit composition[16, 21, 22]. Neto1 maintains the synaptic localization of NR2A subunit-containing NMDA receptors (NMDARs) and thusly mediate long-term potentiation (LTP)[17, 23]. Additionally, Lev9 and Lev10 proteins are responsible for acetylcholine receptor clustering at the neuromuscular junction, thus also regulating synapse composition and function[13]. Future work will pursue the question of whether Csmd2 functions similarly with ionotropic glutamate receptors at the synapse and thusly regulate synapse function. It will be interesting to investigate the physiological consequences of *Csmd2* loss-of-function. Particularly on glutamatergic transmission and whether Csmd2 is required for synapse formation, maturation or function. Given the recent report of a role for Csmd3 in dendrite development[24], it will also be interesting to investigate whether Csmd2 plays a role in elaboration of the dendritic arbor. Future studies focusing on the function of this protein in the central nervous system may lead to a more clear understanding of the molecular mechanisms governing dendrite and synapse formation and function, and possibly the underlying causes of psychiatric disorders associated with defects in neural circuit connectivity, such as schizophrenia and autism spectrum disorder.

## Materials and Methods

### Animals

All experimental methods used in this study were approved by the Institutional Animal Care and Use Committee of the University of Colorado Denver Anschutz Medical Campus and were performed according to approved protocols and guidelines. All experiments involving mouse tissue were conducted using hybrid F1 crosses between the 129X1/SvJ (000691) strain and the C57BL/6J (000664) strain, both obtained from The Jackson Laboratories (JAX).

### Mouse Forebrain Immunohistochemistry

Mouse brains were fixed by transcardial perfusion with 4% paraformaldehyde before dissection and additional fixation for 3 hours at room temperature. Free-floating coronal sections were cut at 75 µm. Prior to immunohistochemical analysis, sections were subjected to antigen retrieval by incubation in 10 mM sodium citrate, pH 6.0 in a pressure cooker set to cook at pressure for 1 minute.

For immunohistochemistry, sections were rinsed with 1x PBS twice for 5 minutes each. Sections were permeabilized with 1x PBS with 0.1% Triton X-100 (Sigma-Aldrich) and then incubated with 10% normal donkey serum (Jackson ImmunoResearch) and 0.1% Triton X-100 in 1x PBS for 1 hour at room temperature. Primary antibodies: Csmd2 (Santa Cruz Biotechnology and Novus Biologicals) 1:200; Olig2 (Millipore) 1:500; Aldhl1 (NeuroMab) 1:500; Ctip2 (Abcam) 1:1000; Satb2 (Abcam) 1:1000; Parvalbumin (Swant) 1:500; Somatostatin (Millipore) 1:250. Sections were incubated in primary antibodies overnight at room temperature. After washing the sections with 1x PBS 3 times for 10 minutes each, relevant AlexaFluor-conjugated secondary antibodies (Life Technologies) were then applied to the sections for 1 hour at room temperature. After rinsing with 1x PBS 3 times for 10 minutes each, 300 nM DAPI (Invitrogen) in 1x PBS was applied for 1 minute. Coverslips were then applied to the sections with ProLong Diamond antifade reagent (Invitrogen). Sections were imaged using a Zeiss LSM 780 confocal microscope.

### Synaptosomal Fractionation from Mouse Forebrain

Preparation of synaptosomal fractions from mouse forebrain homogenate was performed as previously described[18]. Mouse brain homogenates were subjected to separation via ultracentrifugation over a Percoll (GE Healthcare) gradient. The samples obtained from the fractions produced by this protocol were subjected to SDS-PAGE and Western blotting. Crude synaptosomal membranes (P2 pellets) and crude postsynaptic density fractions (TxP) were prepared as previously described[19].

### Western Blot

Protein concentrations were measured with the BCA assay (Pierce, Thermo Scientific) prior to SDS-PAGE and western blot. All protein samples were subjected to SDS-PAGE using 4-15% polyacrylamide gradient gels (Bio-Rad). For cell lysates, 20-30 µg was loaded, while for synaptosomal fraction samples 40 µg of material were applied to the gels. Separated proteins were then electroblotted using a TransBlot Turbo system to TransBlot Turbo Mini-size PVDF membranes (Bio-Rad). Membranes were subsequently blocked with 1x TBS containing 0.1% Tween 20 (1x TBST) with 5% (w/v) blotting-grade blocker (Bio-Rad) and probed with the primary antibody of interest diluted in 1x TBS containing 0.1% Tween 20 and 0.5% blocker at room temperature overnight. Primary antibodies for mouse Csmd2 were used at a dilution of 1:500, PSD95 at 1:1000, and DYKDDDK (FLAG; ThermoFisher Scientific) at 1:500. Membranes were washed 3 times in 1x TBST for 10 minutes each prior to 1 hour of incubation at room temperature with horseradish peroxidase (HRP)-conjugated secondary antibodies used at 1:10,000.

### In Utero *Electroporation*

*In utero* electroporations were performed as described[25]. Briefly, timed pregnant mice (E15.5) were anesthetized and their uterine horns exposed. Endotoxin-free plasmid DNA was injected into the embryos’ lateral ventricles at 1 mg/mL each. For electroporation, 5 pulses separated by 950 ms were applied at 50 V. To target the hippocampus, electrodes were placed in the opposite orientation compared to targeting the neocortex. Embryos were allowed to develop *in utero* and then postnatally until the indicated time.

### Yeast 2-Hybrid Analysis

Yeast two-hybrid screening was performed by Hybrigenics Services, S.A.S., Paris France (http://www.hybrigenics-services.com). The mouse Csmd2 cytoplasmic domain (amino acids 3557-3611) were used as the bait protein and a mouse adult brain cDNA library was the prey.

### 3x FLAG Pull-Down and Co-Immunoprecipitation

For FLAG pull-down co-immunoprecipitation experiments, samples were lysed in a working solution of 50 mM Tris-HCl, 1 mM NaCl, 1% Triton X-100, and 1 mM EDTA, pH 7.6. Every 10 mL of this solution was supplemented with 1 cOmplete ULTRA, Mini, EDTA-free protease inhibitor cocktail tablet (Roche). After lysate preclearing, samples were incubated for 3 hours at 4°C with Anti-DYDDDDK Affinity Gel (Rockland) beads. For all other co-immunoprecipitation experiments, samples were lysed in the aforementioned lysis buffer. After lysate preclearing, samples were incubated with an antibody against the targeted protein of interest overnight at 4°C. After antibody-binding, samples were incubated for 3 hours at 4°C with Protein G Mag Sepharose Xtra (GE Healthcare Life Sciences) beads. After incubation, washes were conducted according to the corresponding manufacturers’ recommended protocol. Samples were eluted from beads via incubation with Laemmli sample buffer (Bio-Rad) at 37°C for 20 minutes prior to analysis by Western blotting.

### Primary Hippocampal Neuron Culture

Primary cultured hippocampal neurons were prepared from embryonic day 17.5 C57Bl/6J mice subjected to *in utero* electroporation, conducted as described. Hippocampal tissue from these mice was manually dissected and dissociated as previously described [26]. 500,000 cells were seeded per well onto poly-D-lysine-coated (Millipore) 12 mm cover slips in 24-well plates in Dulbecco’s Modification of Eagle’s Medium (DMEM; Corning) containing 10% fetal bovine serum (Gibco) and 1% penicillin/streptomycin (Lonza). At 2 DIV, the DMEM-based culture medium was replaced with Minimum Essential Eagle’s Medium (EMEM; Lonza) containing 2.38 mM sodium bicarbonate (Sigma), 2 mM stabilized L-glutamine (Gemini Bio), 0.4% glucose (Sigma). 0.1 mg/mL apo-transferrin (Gemini Bio), 2% Gem21 NeuroPlex Serum-Free Supplement (Gemini Bio), 5% fetal bovine serum (Gibco), and 1% penicillin/streptomycin (Lonza). At 3 DIV, half of the culture medium was replaced with fresh EMEM-based medium. At the indicated times, coverslips were fixed in 4% paraformaldehyde for 20 minutes at room temperature, washed 3x with PBS and mounted onto microscope slides using ProLong Diamond antifade reagent (Invitrogen).

## Acknowledgements

This work was supported by the Children’s Hospital Colorado Program in Pediatric Stem Cell Biology (S.J.F.) and The Boettcher Foundation (S.J.F.). The authors declare no competing financial interests.

